# Biomimetic self-regulation in intrinsically motivated robots

**DOI:** 10.1101/2025.05.11.653305

**Authors:** Oscar Guerrero Rosado, Adrián F. Amil, Ismael T. Freire, Martin Vinck, Paul F.M.J. Verschure

## Abstract

From weaving spiders to hibernating mammals and migratory birds, nature presents numerous examples of organisms exhibiting extraordinary autonomous behaviors that ensure their self-maintenance. However, physiological needs often interact and compete. This requires living organisms to handle them as a coordinated system of internal needs rather than as isolated subsystems. We present an artificial agent equipped with a neural mass model replicating fundamental self-regulatory behaviors observed in desert lizards. Our results demonstrate that this agent not only autonomously regulates its internal temperature by navigating to areas with optimal environmental conditions, but also harmonizes this process with other internal needs, such as energy, hydration, security, and mating. This biomimetic agent outperforms a control agent lacking interoceptive awareness in terms of efficiency, fairness, and stability. Additionally, to demonstrate the flexibility of our framework, we develop a “cautious” agent that prioritizes security over other needs, achieving a Maslow-like hierarchical organization of internal needs. Together, our findings suggest that grounding robot behavior in biological principles of self-regulation provides a robust framework for designing multipurpose, intrinsically motivated agents capable of resolving trade-offs in dynamic environments.

## Introduction

When aiming to build autonomous artificial systems capable of making decisions and behaving without human intervention, a central question arises: If a human is not commanding the robot, where should it ground its decisions? In this paper, we adopt a biomimetic approach to answer this question. We investigate how modeling intrinsic motivation can enhance a more biologically plausible level of autonomy, where goals are grounded in the embodied nature of the agent.

Nature provides numerous examples of organisms that exhibit extraordinary autonomous behaviors. Ranging from weaving spiders to hibernating mammals and migratory birds, these biological systems ground their behavioral strategies on a specific purpose: self-maintenance Maturana and Varela (2012). In order to survive and thrive, autopoietic systems generate autonomous behavior to regulate physiological needs essential for their subsistence Varela et al. (1974); Montévil and Mossio (2015).

Fundamental self-regulatory mechanisms supporting autonomy can be found across organizational scales: from single cells to networks, organs, whole organisms, and groups of organisms Levin (2021). One central self-regulatory mechanism is homeostasis. This self-regulatory mechanism, first described by Walter Cannon Cannon (1939), is defined as the process of maintaining an internal variable within a desired stable range through a negative feedback loop. In this loop, any deviation from this desired range is detected as a homeostatic error. This, in turn, triggers a corrective response from the system.

The concept of homeostasis has not only revolutionized biology but also psychology. One major theory that integrates such self-regulatory mechanisms in the explanation of goal-directed autonomous behavior is the Drive Reduction theory Hull (1952). According to this theory, organisms pursue goals that fulfill internal needs, which are understood in homeostatic terms as deviations from a desired state. The correction of these needs is rewarding for the individuals, encouraging them to reproduce the same behavior in future similar situations Nieh et al. (2016); Grove et al. (2022).

Yet, homeostatic mechanisms fall short when trying to explain self-regulation in its entirety. First, it is challenging to identify steady states in physiological measures. Variables such as heart rate Rajendra Acharya et al. (2006), body temperature Szymusiak (2018), or body mass index Buck and Barnes (1999) exhibit fluctuations over minutes, hours, or days. Second, for autopoietic systems to self-maintain and reproduce, the balance of isolated homeostatic systems is insufficient. Organisms embody a rather complex set of internal needs that often interact and compete with each other, making optimal orchestration critical for survival. Indeed, learning environmental contingencies and developing anticipatory behavior are key to preventing catastrophic deviations from homeostasis. In an attempt to address these issues, Sterling and Eyer Sterling and Eyer (1988) proposed the term *allostasis* as the adaptive stability of the self in response to the demands and opportunities that the environment offers. Thus, allostasis considers stability as a dynamic process that concerns the self rather than its components and occurs in consonance with environmental events.

The hypothalamus, an ancient brain structure found in the brainstem, plays a central role in sensing the physiological states of the individual and orchestrating the derived motivations Stellar (1954). Neurons in specialized hypothalamic nuclei express receptors that track changes in osmolarity, temperature, and energy levels, among others. These nuclei have been reported to hold a relation of mutual inhibition Burnett et al. (2016); Osterhout et al. (2022); Petzold et al. (2023), a competing dynamic that potentially implements a winner-take-all mechanism for motivation orchestration.

Precisely, this competing dynamic based on mutual inhibition has been explored in previous work to reproduce complex animal behavior based on opposing motivations, including security versus exploration and energy versus temperature Rosado et al. (2022). The results showed that a neural mass allostatic model is not only suitable to describe hypothalamic dynamics, but also offers an orchestration mechanism of goal-oriented behavior for both biological and artificial systems. Together with previous biomimetic studies that implement self-regulatory mechanisms Sanchez-Fibla et al. (2010); Vouloutsi et al. (2013); Maffei et al. (2015); Jimenez-Rodriguez et al. (2020), allostasis emerges as a promising theoretical framework for advancing artificial autonomy.

To further evaluate the suitability of neural mass allostatic models for self-regulation in dynamic environments, we turn to a benchmark inspired by natural biology: the Namib desert lizard Houston and McFarland (1976). This ectothermic species exhibits a remarkable behavioral strategy to thermoregulate in extreme conditions. To maintain its body temperature within an optimal range, the lizard alternates between basking in the sun and burrowing into the cooler sand, dynamically adjusting its behavior in response to fluctuating environmental conditions. However, thermoregulation is not its only concern. Pursuing other vital needs, such as hydration, energy acquisition, safety, or mating, often conflicts with thermoregulatory behaviors. As such, the lizard must adaptively orchestrate its actions to maintain multiple internal variables within functional limits, ensuring that no single need is neglected in the long term.

Inspired by self-maintenance behavioral dynamics, this paper proposes an extended neural mass model of allostasis that can manage multiple, and often competing, internal needs. Crucially, unlike previous implementations that relied on abstract simulations (e.g., grid-worlds), the model is deployed and tested within a high-fidelity, dynamic, and multi-agent robotic simulation environment. This setting enables the exploration of embodied self-regulatory behavior under realistic sensory constraints, advancing our understanding of how biologically inspired mechanisms can support artificial autonomy.

## Materials

To replicate the self-regulatory behaviors of the desert lizard in its natural environment, we designed a virtual experimental setup in Webots Michel (2004). This environment was composed of an open arena of 3,5 × 3,5 meters where two static resources representing energy and hydration sources were placed in the top corners. We modeled desert temperature as an oscillating gradient field that originated at the bottom of the arena. This gradient wax and wanes, covering a dynamic portion of the arena along five day-night cycles. As in natural environments, the spatial distribution of resources compels the agent to prioritize which internal need to address. Finally, three e-puck mobile robots represented an allostatic agent (marked in green), a mate agent (marked in blue), and a predator agent (marked in red). While the navigation of the allostatic agent was governed by its internal state, the predator and mate agents explored the environment following random navigation (Fig. 1).

**Figure 1:**
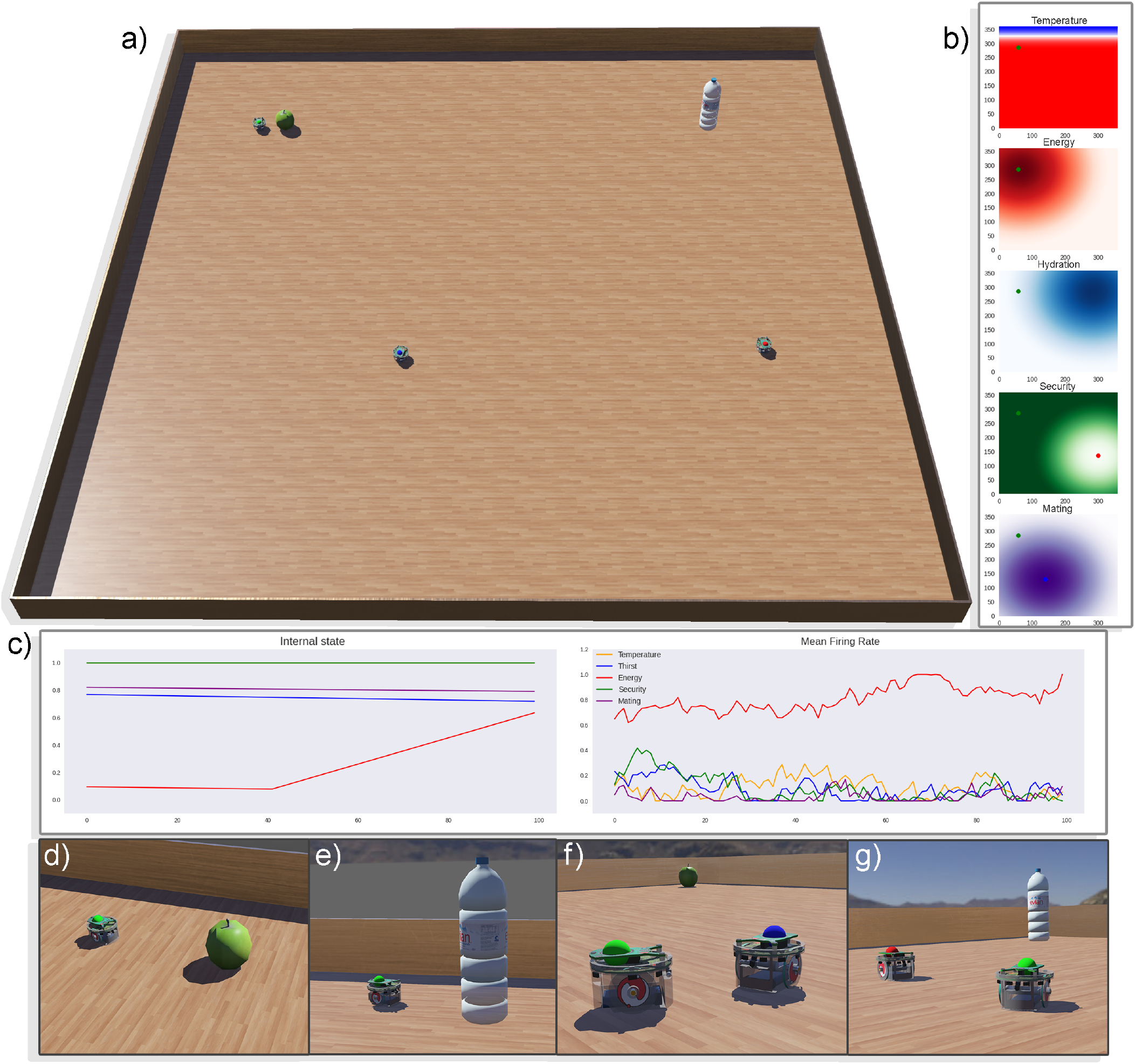
Simulated experimental setup in Webots. a) Top view of the experimental setup, where the e-puck agent (marked in green) is approaching the energy source. b) Panel plotting each gradient field representing the need-fulfilling objects’ location. A green dot marks the location of the allostatic agent. At this precise moment, the allostatic agent is at the top of the energy, temperature, and security gradients. c) The left panel shows the agent’s internal state. Notice that energy levels (red signal) are increasing as the agent is at the food location. The right panel shows the neural mass allostatic model dynamics. At this point, robot navigation is governed by a motivation to restore energy levels. This motivation aligns with the agent’s internal state, as it represents the most pressing need. d) Allostatic agent restoring energy levels by approaching the energy source. e) Allostatic agent restoring hydration levels by approaching the hydration source. f) Allostatic agent fulfilling mating need by approaching its mate (marked in blue). g) Allostatic agent ensuring its security by escaping from a predator (marked in red).

In order to enable goal-oriented navigation without the need to develop cognitive maps, the allostatic agent partially observed overlapping gradient fields that signaled the location of each need-fulfilling resource. The implementation of such a gradient is justifiable as organisms also find in nature physical gradients, such as those based on chemical concentration Stock and Baker (2009) or environmental temperature Houston and McFarland (1976), that they can use to inform their decisions. Moreover, policies generated through reinforcement learning also approximate gradient fields, where states close to rewards augment their state value function Sutton et al. (1998). It has been shown that cognitive systems can also learn and embody such reward-informing gradients as hippocampal place cells cluster near reward locations Hollup et al. (2001), and orbitofrontal cells tend to increase their firing activity as the animal approaches reward Basu et al. (2021). Finally, similar experimental setups have reported gradient-based navigation as an effective approach for robot autonomous navigation Fibla et al. (2010); Sanchez-Fibla et al. (2010); Rosado et al. (2022).

## Methods

A set of five internal states is incorporated into the agent. These states follow homeostatic dynamics; that is, they slowly and continuously decay during periods of inaccessibility to the related resource. Once the agent is in the vicinity (peak area of the gradient field) of a resource, its consumption is considered available, and the internal state is restored to satiation levels. Need magnitudes are defined as homeostatic errors (i.e., Euclidean distance between the current internal state and the homeostatic setpoint). In this study, all setpoints were set at their maximum, encouraging the agent to maximize its internal state.

Homeostatic dynamics differ for each internal state in their leak rate. The temperature and hydration states decayed faster, followed first by energy and then by mating, the latter being the slowest decaying state. Importantly, the security state did not leak. Instead, security levels depended solely on the distance to the predator agent.

The allostatic agent embodied a neural mass model that receives homeostatic errors as inputs and returns motivational drive signals to govern navigation (Fig. 2). This model is grounded in hypothalamic dynamics Burnett et al. (2016); Osterhout et al. (2022); Petzold et al. (2023), where internal drives compete for dominance. Specifically, the model builds on modified Wilson-Cowan equations Wilson and Cowan (1972), as formulated in Amil and Verschure (2021); Rosado et al. (2022), and incorporates both mutual and shared inhibitory projections to regulate competition between drives. The resulting population dynamics are described by the following set of differential equations:

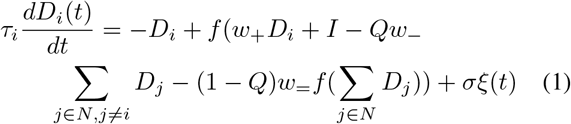

where *f*(*x*) is the logistic *f* − *I* function,

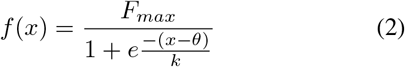

**Figure 2:**
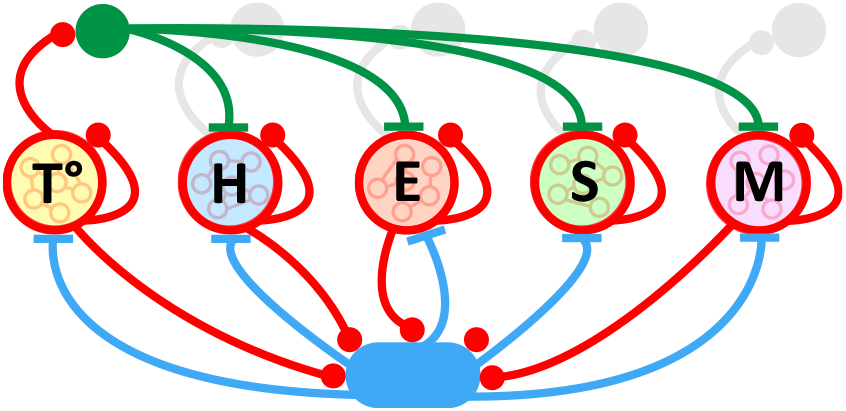
Neural mass multiattractor allostatic model. Our model uses homeostatic errors as excitatory inputs for five state-representing excitatory neural pools: Temperature, Hydration, Energy, Security, and Mating. Each of these populations projects to three targets. First, they maintain their activity over time through recurrent connections. Second, they project to interneurons implementing mutual inhibition (green). Third, they project to interneurons implementing shared inhibition (blue). For illustration purposes, only the temperature neural population illustrates mutual inhibitory connections, but the same topology holds for the other state-representing neural pools.

In these equations, *τ*_*i*_ is the time constant that determines the time scale of the population dynamics. *D* is the set of drive magnitudes, and *D*_*i*_ is the specific motivational drive that is being updated. *I* is the input that each excitatory population receives (that is, homeostatic errors). *w*_+_, *w*_*−*_, and *w*_=_ are the weights for recurrent connections within the excitatory population, mutual inhibition, and shared inhibition, respectively. *Q* represents the mutual/shared inhibition ratio, a variable that controls the level of competition between populations. *σ* is the variance of the Gaussian white noise process represented by *ξ* which follows a normal distribution N(0,1). Finally, *F*_*max*_, *k*, and *θ* are the maximum firing rate, gain, and threshold parameters of the *f − I* logistic curve, respectively (equation 2).

Due to mutual inhibition and sustained homeostatic drives, the model exhibits steady-state dynamics characterized by a system of fixed-point attractors. In this regime, the neural population receiving the largest homeostatic error tends to become dominant, suppressing the activity of competing populations. This dominance persists until the agent’s internal state changes (i.e., until the corresponding homeostatic error is minimized or resolved). The activation of a specific neural population reflects the dominance of a particular drive, which in turn determines the weighting of the associated gradient field used for navigation. As previously described, orientation and internally driven navigation are achieved through local observation of these gradients. For a detailed account of the navigational mechanisms, see Rosado et al. (2022).

Additionally, our model allows for prioritizing the self-regulation of certain internal states by increasing neural pool sensitivity to certain homeostatic errors. In this manner, internal states can be organized similarly to a Maslow hierarchy of needs Maslow (1943). By increasing the model’s sensitivity to security, we developed a cautious agent that prioritized predation avoidance. To validate the performance of the allostatic agent, a third control agent acted as a baseline. Navigation of the control agent was not governed by allostatic dynamics (though it embodied internal states as the others) but by random exploration enhanced with obstacle avoidance.

## Results

The following results are extracted from 20 experimental sessions for each agent condition (allostatic, cautious, and control). Each experimental session simulated five day–night cycles, which, in real time, took approximately 33 minutes to run on the simulation platform.

When analyzing agents’ navigational patterns, we identified a self-organization of trajectories in the allostatic agent. Navigation of this agent was characterized by circular trajectories in the surroundings of the energy and hydration resources and extensive travels covering the entire arena, including the bottom limit where the temperature gradient field is anchored (Fig. 3a and b). This navigational pattern resulted in a clear bias toward the upper part of the arena. As expected, the control agent performing random navigation did not show any sign of navigational bias, with the mean X and Y position located at the center of the arena (Fig. 3a). Occupancy maps revealed goal-oriented self-organized navigation in the allostatic agent, characterized by a marked tendency to explore areas surrounding energy and hydration resources (Fig. 3b). This was not the case of the control agent since there was no clear sign of spatial preference beyond the behavior resulting from obstacle avoidance in the corners of the arena (Fig. 3b). Finally, the cautious agent trajectory and occupancy maps matched the navigational patterns of the allostatic agent; for this reason and to avoid redundancy, these results are not shown.

**Figure 3:**
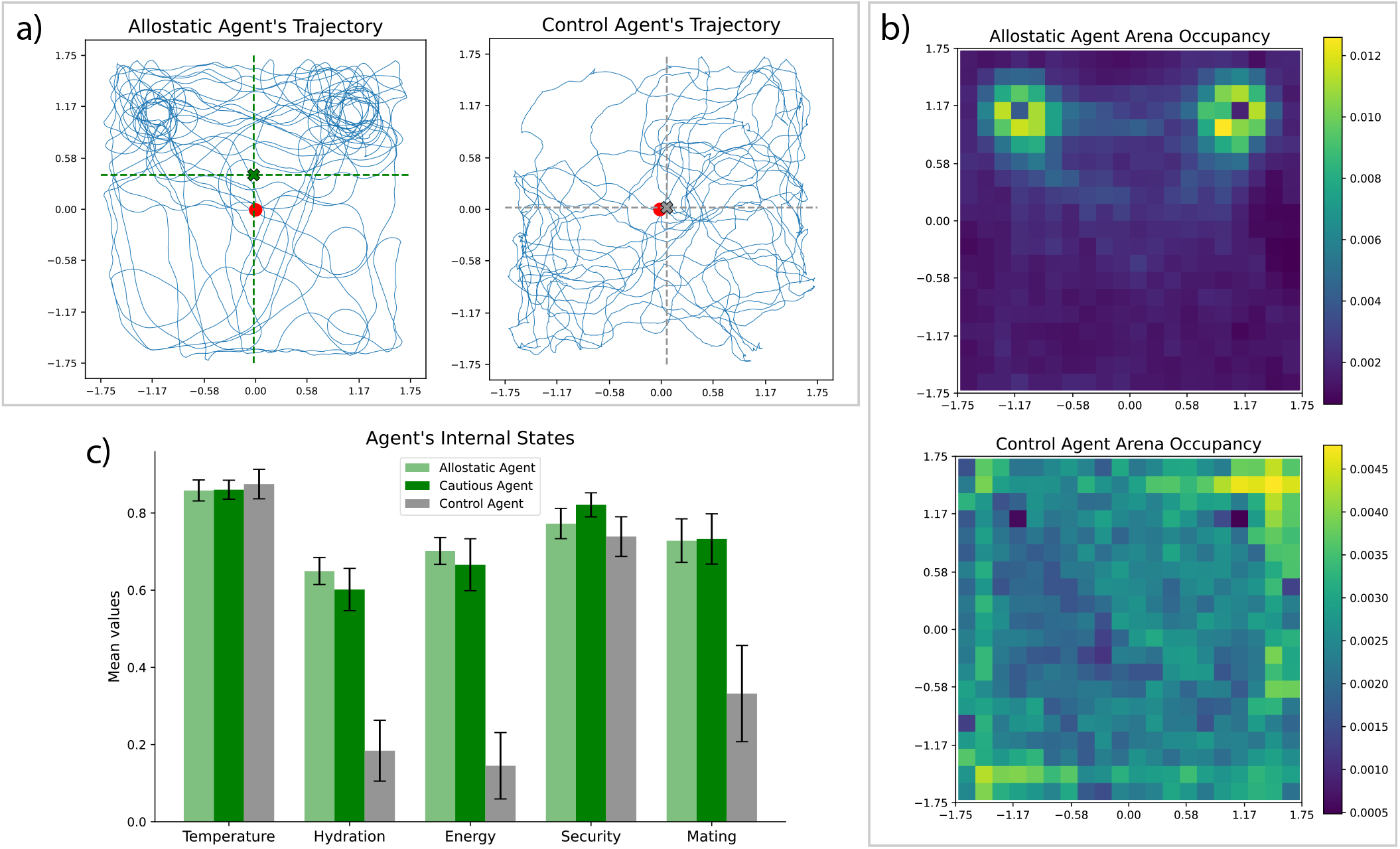
Self-organized navigational patterns allow internal state maintenance. a) Allostatic (left) and control (right) agents’ trajectories. Red dots at the center of the arena represent the agents’ starting positions. Horizontal and vertical dashed lines represent the X and Y-axis mean position, respectively, across 20 experimental sessions recorded with each agent condition (five day-night periods each session). Cross marks represent the mean position of each agent condition (green: allostatic and gray: control). b) Allostatic (top) and control (bottom) agents’ occupancy maps. Colormaps represent the normalized time period spent at each spatial bin across 20 experimental sessions in each agent condition. c) Mean internal state of allostatic (clear green), cautious (dark green), and control (gray) agents across 20 experimental sessions. Error bars indicate standard deviation.

The adoption of a goal-oriented navigation also has an effect on the internal state of the agent. While the control agent fails to regulate its hydration, energy, and mating needs, both allostatic and cautious agents successfully balance these internal drives without compromising thermoregulation or security (Fig. 3c). One may wonder how the control agent maintained high levels of temperature and security. This is because both states are often satisfied across extensive regions of the arena. In fact, the allostatic agent similarly maintained high levels of temperature and security even when it had to travel to locations where these states are compromised. For instance, to balance energy or hydration, the agent travels to the upper part of the arena where temperature often decays, exposing itself to a potential predator.

Moreover, the temporal evolution of the allostatic agent’s internal states reflects the attractor dynamics imposed by the underlying neural mass model. The analysis in this section focuses on energy and hydration, as these states are primarily regulated by internal mechanisms and are not directly influenced by environmental variables such as day–night cycles or the presence of other agents. As designed, the model triggers increased neural firing in the motivational pool associated with an internal state when that state becomes deficient (Fig. 4a). This neural activation drives goal-directed behavior, leading to the replenishment of the corresponding internal need. Once the homeostatic error is resolved, motivational dominance shifts, and another internal state becomes prioritized, thereby suppressing the previously active drive. The attractor landscape reveals that high values in one internal state (for example, hydration or energy) are typically associated with decreasing trends in the other, indicating a trade-off relationship between regulatory priorities (Fig. 4b). This is especially evident in the saddle-like structure in the upper left region, demonstrating winner-takes-all dynamics: When one drive dominates, the competing drives are suppressed. Furthermore, the presence of high-density convergence zones reflects stable attractor regions toward which internal state trajectories consistently flow, supporting the existence of robust, dynamic regulation patterns. The temporal structure of energy and hydration dynamics further reflects their slow, homeostatically driven regulation. Autocorrelation analysis showed that both signals exhibit strong temporal structure at short lags, with high initial autocorrelation that decays within the first few thousand timesteps (Fig. 4c). Beyond this point, the signals fluctuate around zero, indicating low long-range correlation and the absence of persistent drift. These dynamics suggest that internal states are periodically reset through regulatory behavior. Complementing this, spectral analysis using Welch’s method revealed that both energy and hydration signals are dominated by low-frequency components, with most of the power concentrated below 0.05 Hz (Fig. 4d). This supports the interpretation that internal regulation operates on a slow, biologically plausible timescale, consistent with intrinsic motivational cycles shaped by the neural dynamics.

**Figure 4:**
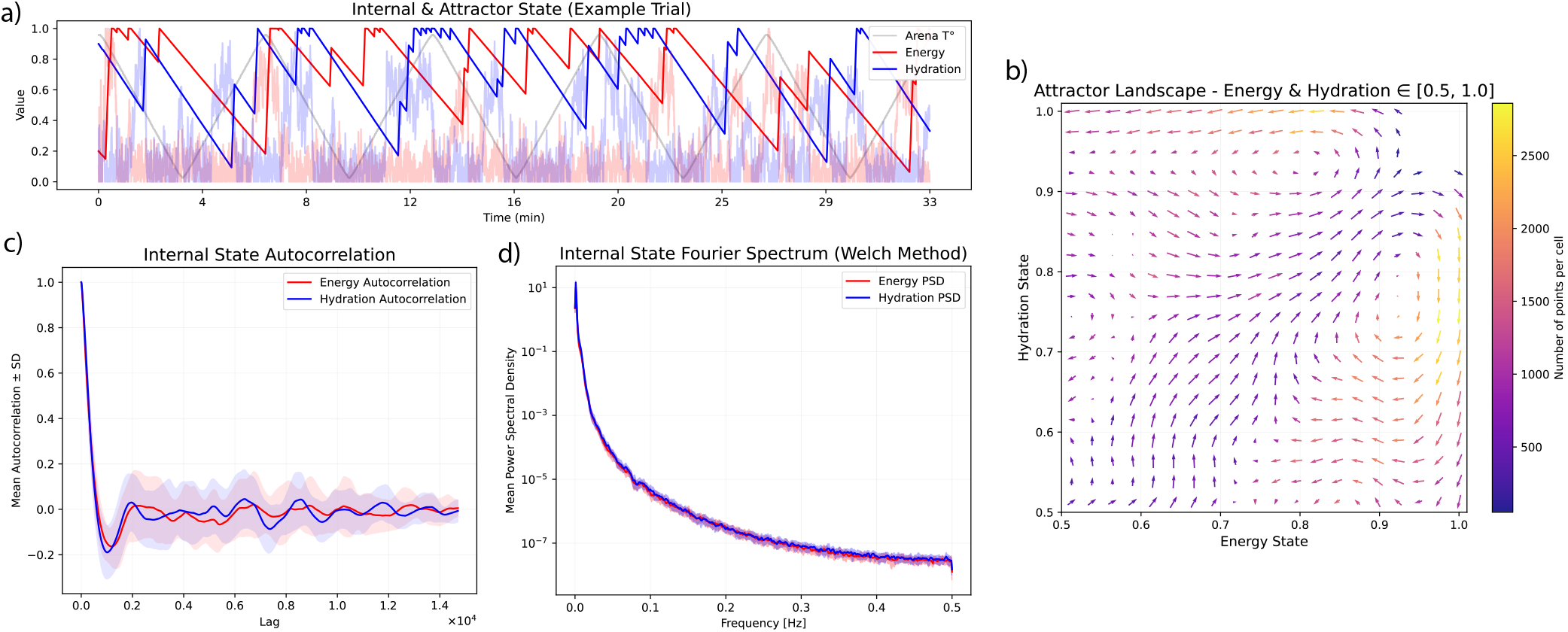
Attractor dynamics in the neural mass allostatic model. a) Time series from a single trial showing the evolution of energy and hydration internal states (solid lines) and the corresponding model outputs (transparent lines). The gray line represents the environmental temperature driven by day–night cycles. b) Attractor landscape in the Energy–Hydration state space, showing average directional flows (arrows) and state density (color-coded). c) Autocorrelation functions of energy and hydration across trials, with shaded areas indicating ±1 standard deviation. d) Power spectral density (PSD) of energy and hydration signals estimated using Welch’s method, highlighting dominant low-frequency dynamics.

To better understand the advantages of adopting an allostatic approach in achieving biological-like autonomy, we computed the game-theoretic metrics that considered the five internal states as a unified system Binmore (2005); Freire et al. (2020). The first metric, *Efficiency*, was computed as:

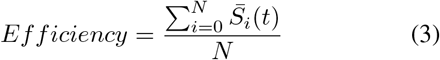

where 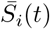 is the internal state *i* at time *t* and *N* is the number of internal states the agent embodies, five in our case. The efficiency metric informs us about the agent’s overall ability to maintain the set of internal states close to their setpoints.

The second metric, *Fairness*, was computed as:

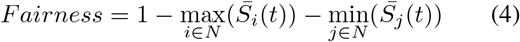

where *max* and *min* 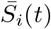 are the maximum and minimum internal states at time *t*. The fairness metric informs us about the agent’s ability to attend to each drive without neglecting any of them in the long term.

The third metric, *Stability*, was computed as the mean square error between the internal states and their setpoints:

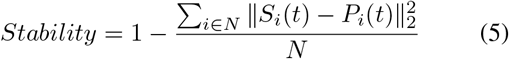

where *S*_*i*_(*t*) is the internal state *i* at time *t* and *P*_*i*_(*t*) is the setpoint for such an internal state at the same moment. The stability metric provides us with a measure of need regulation performance over time.

Additionally, to better understand how these metrics depend on the dynamics of the environmental temperature (i.e., day-night cycles), we grouped data points into bins of environmental temperature and averaged across timesteps and experimental sessions. As a result, we observe that the allostatic agent outperformed the control agent in the three metrics of efficiency, fairness, and stability (Fig. 5a, b, and c). Importantly, besides demonstrating allostasis, correlation values between the game-theoretic measures and the environmental temperature tended to decrease for the allostatic agent. This tendency captures the allostatic agent’s capabilities to reduce environmental influences. It is worth noting that the control agent’s fairness negatively correlated with environmental temperature. This can be explained by the fact that this agent was incapable of self-regulating energy and hydration states, and internal temperature drops approximating energy and hydration levels during nighttime. Thus, when environmental temperature increases, it facilitates thermoregulation and increases the difference between maximum and minimum internal states, in turn decreasing fairness. For the cautious agent, metrics of efficiency, fairness, and stability resembled those of the allostatic agent. To avoid redundancy, these results are not shown in Fig. 5.

**Figure 5:**
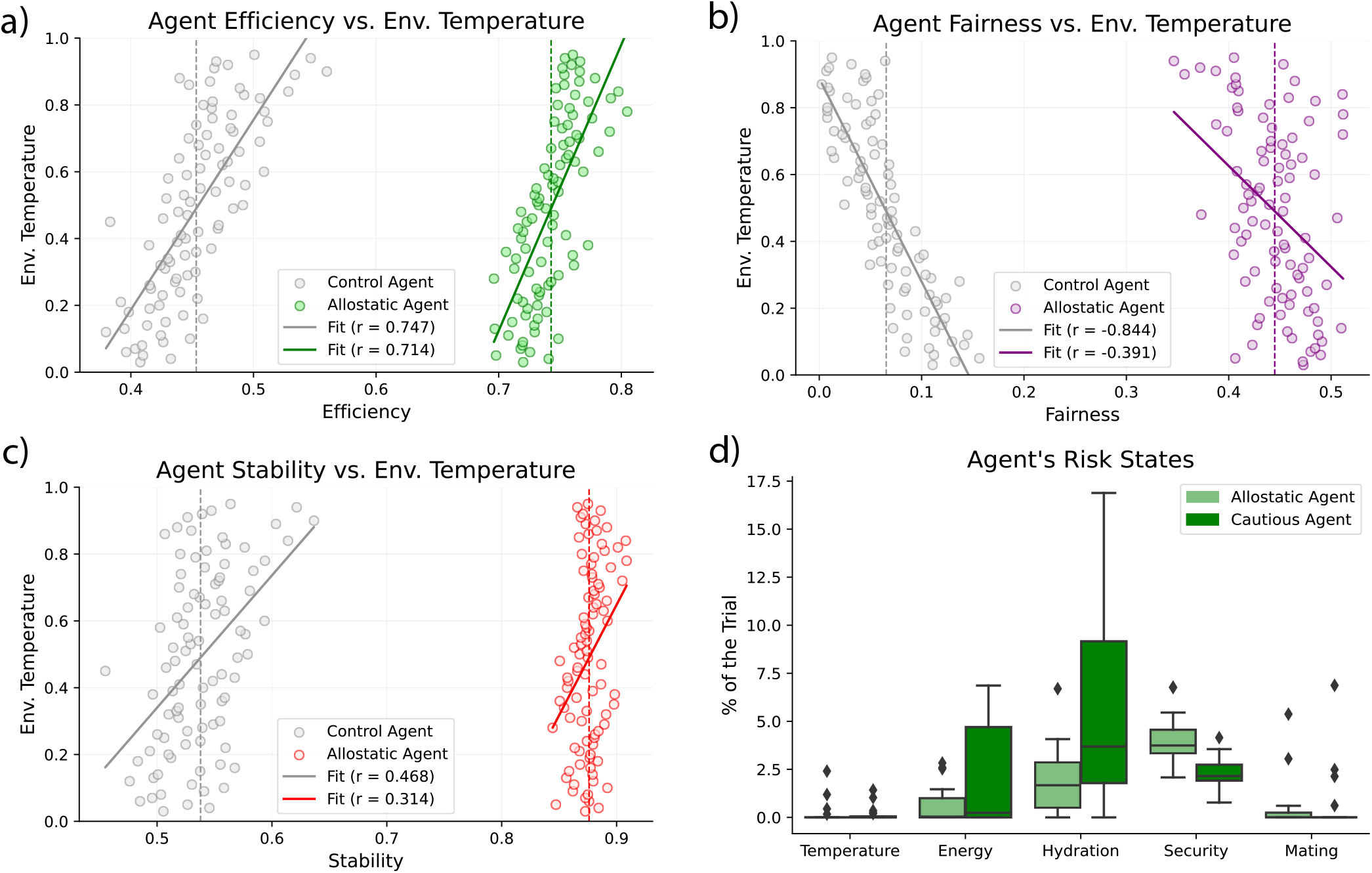
Analysis of game theoretic metrics and periods of risk. a) Efficiency metrics for control (gray) and allostatic (green) agents correlated with environmental temperature. b) Fairness metrics for control (gray) and allostatic (purple) agents correlated with environmental temperature. c) Stability metrics for control (gray) and allostatic (red) agents correlated with environmental temperature. Dashed lines indicate the mean value of the metric across the entire session. Solid lines indicate best-fit regression lines. d) Box plot informing about the period of time both allostatic and cautious agents spent in a state of risk for each of the five internal needs. An internal state was considered at risk when it did not reach the 10% of the setpoint.

Finally, we analyzed the benefits of implementing a hierarchy of needs that prioritizes security. Unlike other needs, which decayed gradually, our experimental setup caused abrupt changes in the security state due to the predator’s random movements. These rapid dynamics posed a significant challenge to the model’s adaptability and occasionally resulted in predation during the allostatic agent’s trials. In fact, for the allostatic agent, security was the internal state that underwent a larger period of the trial under a level of risk (i.e, internal state below 10% of the set point) (Fig. 5d). This pattern changed when the agent’s sensitivity to security-related homeostatic errors was increased. The more cautious agent showed a greater ability to avoid predation by reducing the time spent under threat. However, prioritizing security came at a cost: it led to longer periods in which the internal states for hydration and energy remained in a state of risk (Fig. 5d).

## Discussion

In this study, we propose that robot autonomy can be enhanced by integrating core principles of biological autonomy, enabling artificial agents to self-regulate, adapt, and survive in complex environments. To test this hypothesis, we conducted an experiment in which a robot was tasked with self-regulating multiple internal needs in a setting designed to replicate desert environmental conditions. For the artificial agent to self-regulate, we designed and integrated a neural mass model of allostasis using a hypothalamic neural architecture as design constraints. We observe that an allostatic agent demonstrates a remarkable level of autonomy, (1) orchestrating its internal states so that none of them are neglected, (2) minimizing the periods where a need may represent a survival risk, (3) adapting to the dynamics of the environment and of the other agents, and (4) becoming cautious when a Maslow-like hierarchy of needs prioritizes security. Using game-theoretic measures, we also observed that the allostatic agent outperforms baseline performance while becoming more resistant to environmental temperature fluctuations.

Together, these results support the idea that adopting a biomimetic approach toward self-regulation can significantly enhance autonomy in artificial systems. A potential industrial implementation could involve a robot builder deployed on the Moon to construct base camps. In such an environment, the agent must thermoregulate, as extreme lunar temperatures can damage critical components such as batteries or sensors. Simultaneously, the robot must coordinate multiple competing tasks: It needs to recharge its batteries by orienting its solar panels towards sunlight, forage for construction materials such as lunar regolith, collaborating with other agents to achieve goals unavailable to the individual, and maintaining safety by avoiding high-radiation areas, unstable terrain, or zones with elevated meteorite risk. Equipping the builder with the allostatic model described here could enhance its autonomy, reducing reliance on human teleoperation.

Nonetheless, such an industrial implementation is challenged by a key limitation of our study. As described, robot navigation relied on predefined gradient fields provided by the experimenter. To navigate, autonomous robots need to learn their own cognitive maps flexibly and reliably. Recent work has demonstrated that a hippocampal-inspired model can generate motivationally informed cognitive maps Guerrero Rosado et al. (2025), as observed in animal models Kennedy and Shapiro (2009). This methodology has proven to be effective for learning optimal policies toward distinct objectives where reward values depend on the motivational state of the agent. Future research aims to integrate both biologically grounded models to support lifelong learning in intrinsically motivated agents.

Another path to explore is the extension of the hierarchy of needs to account for more abstract internal needs such as self-esteem, status, and self-actualization. How those needs emerge in cognitively complex organisms remains an open question. One possibility, inspired by constraint closure Montévil and Mossio (2015); Wilson and Prescott (2022), is that they emerge to support basic needs. In this way, belonging could ensure the availability of resources by sharing within a community. Social rank could increase the resource allocation within that community, and self-actualization could assist social rank.

In conclusion, our research provides evidence for the advantage of grounding artificial autonomy in biological principles. By doing so, we demonstrate that artificial systems can become efficient, adaptive, and robust in scenarios where their survival depends on their self-maintenance capabilities.

## Acknowledgements

This study was funded by the European Union as part of Counterfactual Assessment and Valuation for Awareness Architecture—CAVAA (European Innovation Council (EIC), 101071178).

Code available at: https://github.com/SRU-CAVAA/WP1-Desert_lizard

## References

Amil, A. F. and Verschure, P. F. (2021). Supercritical dynamics at the edge-of-chaos underlies optimal decision-making. Journal of Physics: Complexity, 2(4):045017.

Basu, R., Gebauer, R., Herfurth, T., Kolb, S., Golipour, Z., Tchumatchenko, T., and Ito, H. T. (2021). The orbitofrontal cortex maps future navigational goals. Nature, 599(7885):449–452.

Binmore, K. (2005). Natural justice. Oxford university press.

Buck, C. L. and Barnes, B. M. (1999). Annual cycle of body composition and hibernation in free-living arctic ground squirrels. Journal of Mammalogy, 80(2):430–442.

Burnett, C. J., Li, C., Webber, E., Tsaousidou, E., Xue, S. Y., Brüning, J. C., and Krashes, M. J. (2016). Hunger-driven motivational state competition. Neuron, 92(1):187–201.

Cannon, W. B. (1939). The wisdom of the body.

Fibla, M. S., Bernardet, U., and Verschure, P. F. (2010). Allostatic control for robot behaviour regulation: An extension to path planning. In 2010 IEEE/RSJ International Conference on Intelligent Robots and Systems, pages 1935–1942. IEEE.

Freire, I. T., Moulin-Frier, C., Sanchez-Fibla, M., Arsiwalla, X. D., and Verschure, P. F. (2020). Modeling the formation of social conventions from embodied real-time interactions. PloS one, 15(6):e0234434.

Grove, J. C., Gray, L. A., La Santa Medina, N., Sivakumar, N., Ahn, J. S., Corpuz, T. V., Berke, J. D., Kreitzer, A. C., and Knight, Z. A. (2022). Dopamine subsystems that track internal states. Nature, 608(7922):374–380.

Guerrero Rosado, O., Amil, A. F., Freire, I. T., Vinck, M., and Verschure, P. F. (2025). Motivational cognitive maps for self-regulated autonomous navigation. bioRxiv, pages 2025–03.

Hollup, S. A., Molden, S., Donnett, J. G., Moser, M.-B., and Moser, E. I. (2001). Accumulation of hippocampal place fields at the goal location in an annular watermaze task. Journal of Neuroscience, 21(5):1635–1644.

Houston, A. and McFarland, D. (1976). On the measurement of motivational variables. Animal Behaviour, 24(2):459–475.

Hull, C. L. (1952). A behavior system; an introduction to behavior theory concerning the individual organism.

Jimenez-Rodriguez, A., Prescott, T. J., Schmidt, R., and Wilson, S. (2020). A framework for resolving motivational conflict via attractor dynamics. In Conference on biomimetic and biohybrid systems, pages 192–203. Springer.

Kennedy, P. J. and Shapiro, M. L. (2009). Motivational states activate distinct hippocampal representations to guide goal-directed behaviors. Proceedings of the National Academy of Sciences, 106(26):10805–10810.

Levin, M. (2021). Life, death, and self: Fundamental questions of primitive cognition viewed through the lens of body plasticity and synthetic organisms. Biochemical and Biophysical Research Communications, 564:114–133.

Maffei, G., Santos-Pata, D., Marcos, E., Sánchez-Fibla, M., and Verschure, P. F. (2015). An embodied biologically constrained model of foraging: from classical and operant conditioning to adaptive real-world behavior in dac-x. Neural Networks, 72:88–108.

Maslow, A. H. (1943). A theory of human motivation. Psychological review, 50(4):370.

Maturana, H. R. and Varela, F. J. (2012). Autopoiesis and cognition: The realization of the living, volume 42. Springer Science & Business Media.

Michel, O. (2004). Cyberbotics ltd. webots™: professional mobile robot simulation. International Journal of Advanced Robotic Systems, 1(1):5.

Montévil, M. and Mossio, M. (2015). Biological organisation as closure of constraints. Journal of theoretical biology, 372:179–191.

Nieh, E. H., Vander Weele, C. M., Matthews, G. A., Presbrey, K. N., Wichmann, R., Leppla, C. A., Izadmehr, E. M., and Tye, K. M. (2016). Inhibitory input from the lateral hypothalamus to the ventral tegmental area disinhibits dopamine neurons and promotes behavioral activation. Neuron, 90(6):1286–1298.

Osterhout, J. A., Kapoor, V., Eichhorn, S. W., Vaughn, E., Moore, J. D., Liu, D., Lee, D., DeNardo, L. A., Luo, L., Zhuang, X., et al. (2022). A preoptic neuronal population controls fever and appetite during sickness. Nature, 606(7916):937–944.

Petzold, A., Van den Munkhof, H. E., Figge-Schlensok, R., and Korotkova, T. (2023). Complementary lateral hypothalamic populations resist hunger pressure to balance nutritional and social needs. Cell Metabolism, 35(3):456–471.

Rajendra Acharya, U., Paul Joseph, K., Kannathal, N., Lim, C. M., and Suri, J. S. (2006). Heart rate variability: a review. Medical and biological engineering and computing, 44:1031–1051.

Rosado, O. G., Amil, A. F., Freire, I. T., and Verschure, P. F. (2022). Drive competition underlies effective allostatic orchestration. Frontiers in Robotics and AI, 9:1052998.

Sanchez-Fibla, M., Bernardet, U., Wasserman, E., Pelc, T., Mintz, M., Jackson, J. C., Lansink, C., Pennartz, C., and Verschure, P. F. (2010). Allostatic control for robot behavior regulation: a comparative rodent-robot study. Advances in Complex Systems, 13(03):377–403.

Stellar, E. (1954). The physiology of motivation. Psychological review, 61(1):5.

Sterling, P. and Eyer, J. (1988). Allostasis: A new paradigm to explain arousal pathology. In Fisher, S.and Reason, J., editors, Handbook of Life Stress, Cognition and Health, pages 629–649. John Wiley & Sons.

Stock, J. B. and Baker, M. (2009). Chemotaxis. In Encyclopedia of Microbiology, Third Edition, pages 71–78. Elsevier.

Sutton, R. S., Barto, A. G., et al. (1998). Reinforcement learning: An introduction, volume 1. MIT press Cambridge.

Szymusiak, R. (2018). Body temperature and sleep. Handbook of clinical neurology, 156:341–351.

Varela, F. G., Maturana, H. R., and Uribe, R. (1974). Autopoiesis: The organization of living systems, its characterization and a model. Biosystems, 5(4):187–196.

Vouloutsi, V., Lallée, S., and Verschure, P. F. (2013). Modulating behaviors using allostatic control. In Biomimetic and Biohybrid Systems: Second International Conference, Living Machines 2013, London, UK, July 29–August 2, 2013. Proceedings 2, pages 287–298. Springer.

Wilson, H. R. and Cowan, J. D. (1972). Excitatory and inhibitory interactions in localized populations of model neurons. Biophysical journal, 12(1):1–24.

Wilson, S. P. and Prescott, T. J. (2022). Scaffolding layered control architectures through constraint closure: insights into brain evolution and development. Philosophical Transactions of the Royal Society B, 377(1844):20200519.

